# Multi-Modal Deep Learning Integrates Spatial Topologies and Sequential Motifs to Identify Class I HDAC Inhibitors as Pan-Cancer Therapeutics

**DOI:** 10.64898/2026.04.22.720196

**Authors:** Siyuan Tong, Wen Zhang, Shiliang Ji

**Author notes:** Corresponding authors: Shiliang Ji and Siyuan Tong. This research was funded by the National Natural Science Foundation of China (grant no. 823B2095), and the Suzhou Medical Innovation Applied Research (grant nos. SYWD2024255, SYW2025185).

## Abstract

The molecular characterization of human solid tumors has introduced immense genomic complexity and intra-tumoral diversification. Converting these detailed, multi-omic profiles right into workable, broad-spectrum therapeutics continues to be an formidable bottleneck in precision oncology. Traditional computational drug repurposing strategies largely rely on single-modality chemical descriptors, which frequently fail to capture the systemic transcriptomic interactions within the highly dynamic tumor microenvironment. Here, this study presents a robust multi-modal deep learning framework that synergistically integrates two-dimensional (2D) molecular graphs via Graph Neural Networks (GNNs) and chemical functional group patterns via self-attention Transformers. By mapping this dual-stream chemical feature space to the perturbational transcriptomic signatures (LINCS L1000) of 22 distinct cancer types from The Cancer Genome Atlas (TCGA), a vast library of over 28,000 small-molecule compounds was computationally screened. The developed multi-modal architecture achieved state-of-the-art predictive accuracy, significantly outperforming traditional single-modality baseline models. Strikingly, the comprehensive pan-cancer transcriptomic reversal landscape identified a persistent convergence of non-oncology drugs exhibiting potent broad-spectrum anti-tumor potential. Specifically, Class I Histone Deacetylase (HDAC) inhibitorsmost notably TC-H-106, RG2833, and Tianeptinaline, agents originally developed to penetrate the blood-brain barrier for neurodegenerative and psychiatric disordersemerged as top therapeutic candidates across lung adenocarcinoma (LUAD), bladder urothelial carcinoma (BLCA), and rectum adenocarcinoma (READ). Subsequent high-dimensional network pharmacology and functional enrichment analyses confirmed that these agents robustly suppress essential oncogenic pathways, specifically collapsing the G1/S phase transition and DNA damage repair machineries. Furthermore, structural validation via molecular docking and force-field thermodynamics confirmed the highly stable physical binding affinity (Vina score: -7.0 kcal/mol, MMFF94 Energy: 64.76 kcal/mol) of TC-H-106 to the HDAC1 catalytic pocket. Kaplan-Meier survival analysis based on TCGA gene expression stratification underscored the significant prognostic benefit of targeting this epigenetic axis. Collectively, these findings introduce a powerful multi-modal AI framework for systems-level drug repurposing and highlight brain-penetrant Class I HDAC inhibitors as highly promising candidates for pan-cancer epigenetic therapy.

## I. Introduction

Cancer is an inherently complex, polygenic, and highly dynamic disease driven by a multitude of genomic, transcriptomic, and epigenetic alterations [1]. Over the past decade, landmark international initiatives, notably The Cancer Genome Atlas (TCGA) and the Pan-Cancer Analysis of Whole Genomes (PCAWG) consortium, have systematically cataloged these molecular aberrations across thousands of primary and metastatic tumors [2]. These monumental sequencing efforts have fundamentally re-shaped the contemporary understanding of oncology, revealing that while specific oncogenic drivers dictate lineage-dependent tumor evolution, there exist profound shared molecular vulnerabilities, convergent evolutionary trajectories, and overlapping transcriptomic dependencies across disparate tissue origins [3]–[5]. Despite this deep, systems-level mechanistic understanding of tumor biology, the clinical translation of these massive genomic discoveries into effective, broad-spectrum targeted therapeutics has significantly lagged. The traditional de novo drug discovery pipeline is notoriously time-consuming, highly capital-intensive, and fraught with staggering attrition ratesoften exceeding 90% during clinical translationnecessitating radical, paradigm-shifting approaches to oncology drug development [6].

Drug repurposing (or repositioning)the application of established pharmacological agents with known safety and pharmacokinetic profiles to novel therapeutic indicationspresents a highly efficient, cost-effective strategy to bypass the exorbitant costs and extended Phase I safety evaluation phases of traditional drug discovery [7]. The advent of large-scale pharmacogenomic databases, such as the Library of Integrated Network-Based Cellular Signatures (LINCS) L1000 dataset, has catalyzed the computational matching of drug-induced perturbational profiles with disease-specific transcriptomic signatures [8]. The underlying premise of transcriptomic reversal asserts that if a compound induces a gene expression signature that is highly anti-correlated with the pathological disease signature, it holds therapeutic potential to revert the diseased state toward physiological homeostasis [9], [10]. Traditional computational drug repurposing strategies largely rely on simplistic similarity metrics (e.g., cosine similarity, Kolmogorov-Smirnov statistics) or single-modality chemical descriptors [11]–[13], building upon foundational AI breakthroughs in molecular property prediction [14], [15], driven by recent artificial intelligence advancements such as DeepDR [16], MolBERT [17], ChemBERTa [18], and DeepPurpose [19]. Nevertheless, most existing frameworks focus either on molecular structure representation or transcriptomic similarity, rarely integrating both modalities in a unified predictive architecture. Crucially, these existing models often treat molecules as a monolithic feature vector, fundamentally ignoring the complementary dual nature of small molecules as both structural “graphs” and semantic “sequences.” This uni-dimensional limitation fails to capture the intricate 3D spatial architecture, stereochemistry, and localized functional group interactions that strictly dictate a molecule’s true pharmacological behavior in a highly dynamic biological system.

In the past two years (2024-2025), the advent of multi-modal foundation models has revolutionized the life sciences, demonstrating that integrating orthogonal data modalities drastically enhances predictive resolution [20]– [22]. Building upon this paradigm, Graph Neural Networks (GNNs) have emerged as the premier architecture for learning molecular topology, excelling at capturing the *spatial local environment* of atoms and bonds via iterative message passing [23]. Concurrently, Transformer-based architectures have demonstrated exceptional capability in modeling the *global semantic dependencies* of chemical substructures via self-attention mechanisms [24]– [26]. Despite these advancements, the synergistic fusion of these two orthogonal perspectivesleveraging GNNs for local atomic neighborhoods and Transformers for global functional motifsremains a critical untapped frontier in systems-level pan-cancer screening.

To address this critical methodological gap, a novel multi-modal deep learning framework was engineered that synergistically integrates spatial GNNs and sequential self-attention Transformers. By fusing the topological representation of molecular graphs with the localized functional feature space of Morgan fingerprints, the dual-stream architecture accurately maps the expansive chemical space to complex perturbational transcriptomic responses. This advanced computational model was systematically deployed to screen a vast library of over 28,000 small molecules against the highly curated disease signatures of 22 distinct TCGA cancer types. The comprehensive in silico screening unveiled a high-resolution pan-cancer transcriptomic reversal landscape, successfully identifying both established lineage-specific targeted agents and highly unexpected broad-spectrum candidates.

Crucially, the multi-modal analysis revealed a robust and persistent convergence on Class I Histone Deacetylase (HDAC) inhibitors as top-tier therapeutic candidates across anatomically distinct lineages. It is imperative to acknowledge that HDAC inhibitors are established anti-cancer agents [27]–[29]. First-generation HDAC inhibitors, such as vorinostat and romidepsin, have successfully achieved FDA approval for hematological malignancies [30], [31], highlighting the broader pan-cancer potential of epigenetic therapy [32], [33]. However, their clinical translation to solid tumors has been severely hampered by poor pharmacokinetic profiles and an inability to cross the blood-brain barrier (BBB). The non-oncology candidates identified by the proposed AI (e.g., TC-H-106, RG2833) were originally engineered for neurological disorders. Their exceptional BBB permeability offers a paradigm-shifting advantage: brain-penetrant epigenetic modulators capable of simultaneously targeting primary tumors and CNS metastases.

To computationally realize this discovery, the drug re-purposing task is formulated as a high-dimensional regression problem. Formally, the input to the multi-modal framework is a structural and semantic representation of a given compound-cell line pair. The output is a predicted continuous gene perturbation vector corresponding to 12,328 high-fidelity transcriptomic features. The network is optimized end-to-end by minimizing the Mean Squared Error (MSE) between the predicted perturbation vector and the empirically derived LINCS L1000 signature.

## II. Methods

### A. TCGA Pan-Cancer Data Acquisition and Transcriptomic Processing

To establish a comprehensive and statistically robust pan-cancer landscape, bulk RNA-sequencing (RNA-seq) datasets spanning 22 distinct solid tumor lineages were systematically retrieved from The Cancer Genome Atlas (TCGA) via the Genomic Data Commons (GDC) Data Portal [34]. The raw HTSeq read count matrices were aggregated, and low-expression genes across the cohorts were strictly filtered. Normalization was performed utilizing the Trimmed Mean of M-values (TMM) method to account for varying sequencing depths, followed by conversion to log2-transformed counts per million (log2-CPM) using the *voom* transformation [35].

Disease-specific pathological transcriptomic signatures were rigorously derived by computing the differential gene expression (log2 fold-change) between primary tumor tissues and patient-matched or adjacent normal healthy samples. The empirical Bayes moderation of standard errors was applied via the *limma* package. Statistical significance was defined using a stringent false discovery rate (FDR) adjusted *P* -value threshold of *<* 0.05 (Benjamini-Hochberg correction).

### B. LINCS L1000 Pharmacological Dataset Construction

For the pharmacological perturbation feature space, Level 5 signature profiles were procured from the LINCS L1000 dataset (GEO accession: GSE92742) [8]. To ensure rigorous model training and strictly avoid data leakage arising from different doses or identical cell lines treated with the same compound, the sample space was explicitly defined as unique drug-cell line perturbation pairs. Following stringent quality control (filtering out low Transcriptional Activity Score profiles and restricted structures), the final high-fidelity dataset comprised *N* = 55, 695 unique perturbation signatures. The target variable (ground truth) for the deep learning framework was strictly defined as the 12,328-dimensional continuous vector representing the gene expression fold-change.

### C. Chemical Representation and Feature Engineering

A dual-representation strategy was employed to capture both the local topology and global substructural patterns of each compound. Molecular structures, encoded as Simplified Molecular Input Line Entry System (SMILES) strings, were programmatically parsed utilizing the RDKit cheminformatics suite [36].

For topological representation, molecules were converted into 2D undirected graphs 𝒢 = (𝒱, ℰ). Each node *v*_*i*_ ∈ 𝒱 representing an atom was initialized with a highly descriptive feature vector encompassing atomic number, hybridization state, implicit valence, formal charge, and aromaticity flags. Each edge *e*_*ij*_ ∈ ℰ representing a chemical bond was parameterized with bond type (single, double, triple, aromatic) and stereochemical configuration. For functional representation, the SMILES strings were hashed into high-dimensional Extended-Connectivity Fingerprints (ECFP4, specifically Morgan fingerprints with radius = 2, length = 1024 bits), effectively capturing the presence and frequency of localized chemical motifs and pharmacophores.

### D. Spatial Topology Learning via Graph Neural Networks (GNN)

To extract latent spatial embeddings from the molecular graphs, a deep Graph Convolutional Network (GCN) was constructed. The message-passing neural network aggregates neighborhood information iteratively to update node representations [37], [38]. Let 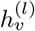 denote the hidden state of node *v* at the *l*-th layer. The graph convolution operation is mathematically defined as:

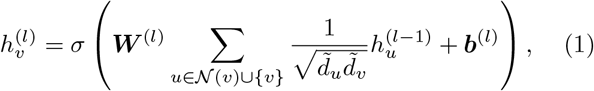

where 𝒩 (*v*) denotes the immediate neighborhood of atom 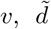 represents the normalized degree of the node to mitigate hub-node dominance, ***W*** ^(*l*)^ and ***b***^(*l*)^ are the layer-specific trainable weight matrix and bias vector, respectively, and *σ* represents the non-linear LeakyReLU activation function. Following successive layers of message passing, a global Readout function (Global Mean Pooling) was applied to aggregate all atom-level embeddings into a singular, fixed-size, permutation-invariant graph topological vector, denoted as ***v***_*graph*_ ∈ ℝ^*d*^.

### E. Sequential Motif Extraction via Self-Attention Transformers

Concurrently, the 1024-bit Morgan fingerprints were processed to capture long-range substructural dependencies using a customized self-attention Transformer encoder [24]. The fingerprint vector was treated as a pseudo-sequence where each bit position was interpreted as a token representing the presence of a specific chemical substructure. The input sequence was projected into an embedding space and processed through multiple multi-head attention (MHA) layers. The scaled dot-product attention mechanism dynamically calculates the relational importance between disparate functional groups:

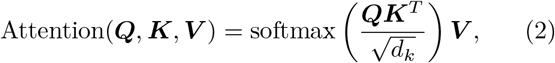

where the Queries (***Q***), Keys (***K***), and Values (***V***) are learned linear projections of the chemical input sequence, and *d*_*k*_ is the scaling factor denoting the dimensionality of the keys. The outputs from multiple attention heads were concatenated, passed through a Position-wise Feed-Forward Network (FFN) with Layer Normalization, and collapsed to generate the dense functional feature vector, ***v***_*func*_ ∈ ℝ^*d*^.

### F. Cross-Modal Attention Fusion and Rigorous Split Strategy

To fuse the modalities, a Cross-Modal Attention mechanism was engineered, allowing the spatial graph topology to actively query the sequential functional representations. Mathematically, this is defined as:

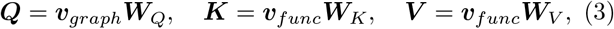

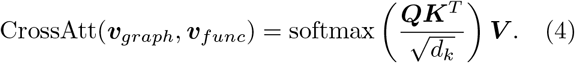

This advanced fusion ensures that the model dynamically emphasizes specific global functional groups based on their precise local spatial atomic context. The fused representation, encompassing both structural physics and chemical semantics, was propagated through a Multi-Layer Perceptron (MLP) decoding head. The network was trained end-to-end utilizing the AdamW optimizer to minimize the Mean Squared Error (MSE) loss between the predicted perturbation vector and the empirical LINCS signature. The explicit model hyperparameter configurations governing the neural network architecture and training dynamics are detailed in Table I.

**TABLE I:** Hyperparameter Configuration for the Dual-Stream Multi-Modal Architecture.

| Module | Hyperparameter | Value |
| --- | --- | --- |
| Graph Neural Network (GNN) | Number of Layers | 3 |
| | Hidden Size ($d$) | 512 |
|  | Attention Heads | 8 |
| Functional Fingerprint (MLP) | Input Dimension | 1024 bits |
| | Hidden Size ($d$) | 1024 |
|  | Dropout Rate | 0.3 |
| Cross-Modal Fusion Layer | Intermediate Size | 2048 |
|  | Dropout Rate | 0.4 |
| Training Configuration | Batch Size | 128 |
|  | Max Epochs | 60 |
| | Learning Rate (AdamW) | $2 \times 10^{-4}$ |

To rigorously assess true generalizability and definitively prevent data leakage, a strict “Drug-Cell Line Pair” split strategy was implemented. Samples were clustered based on their unique compound-cell line combinations, and a Group Shuffle Split was performed (85% training, 15% testing). This mandate ensures that the specific interaction between a given drug and a specific cellular microenvironment in the test set is entirely unseen during training, mirroring the real-world scenario of repurposing known drugs into novel oncological contexts.

### G. Baseline Models for Performance Benchmarking

To rigorously evaluate the proposed architecture, it was benchmarked against traditional single-modality baseline models, including functional fingerprint architectures analogous to DeepPurpose [19] and AttentiveFP [39], as well as specialized transcriptomic prediction frameworks inspired by DeepCE [40] and ChemCPA [41]. To ensure a rigorously fair, apples-to-apples performance comparison, all baseline architectures were re-implemented and trained from scratch on the standardized LINCS L1000 dataset, rather than relying on disparate pre-trained weights. Performance was strictly evaluated using multi-dimensional regression metrics: Mean Squared Error (MSE), Mean Absolute Error (MAE), Pearson correlation (*R*), and Spearman rank correlation (*ρ*).

### H. Weighted Transcriptomic Reversal Scoring (wTRS) Framework

The therapeutic aptitude of each compound against specific cancer types was systematically quantified utilizing a novel Weighted Transcriptomic Reversal Score (wTRS). Recognizing that not all 978 LINCS landmark genes contribute equally to the oncogenic phenotype, a dynamic weighting factor was introduced (*ω*_*g*_), defined as the absolute log2 fold-change magnitude (|*D*_*g*_|) of the gene in the TCGA primary tumor cohort. Genes exhibiting extreme dysregulation in cancer are thus assigned strictly higher pharmacological priority. Let ***D*** represent the disease signature vector and ***P*** represent the predicted drug perturbation vector. The capacity of a drug to act as a therapeutic antagonist is mathematically defined as:

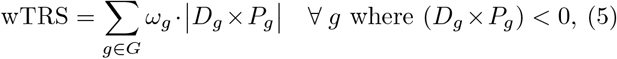

where *ω*_*g*_ = |*D*_*g*_|. By amplifying the contribution of highly perturbed oncogenes and filtering out positively correlated responses, the wTRS precisely quantifies the drug’s capacity to forcefully revert the pathological transcriptomic landscape.

### I. Topological Network Pharmacology and Functional Enrichment

To elucidate the complex mechanisms of action for top-tier candidates (e.g., TC-H-106), a network pharmacology pipeline was implemented. Putative regulatory targets were identified by isolating the genes driving the highest local reversal scores. These targets were mapped onto the high-confidence human protein-protein interaction (PPI) interactome (STRING v11.5). The resulting bipartite drug-target-disease networks were visualized and analyzed using Cytoscape (v3.9.1) [42]. Biological mechanisms were deconvoluted using Gene Ontology (GO) biological processes and Kyoto Encyclopedia of Genes and Genomes (KEGG) pathway databases utilizing established analytical packages [43], [44]. Enrichment was computed using a hypergeometric distribution test, with the statistical significance stringently controlled via Benjamini-Hochberg FDR correction (*P*_*adj*_ < 0.05).

### J. Clinical Survival Analysis and Cox Proportional Hazards Modeling

To bridge computational predictions with in vivo clinical prognosis, survival analyses were executed using corresponding patient clinical metadata from TCGA. Patient cohorts were stratified into discrete high-risk and low-risk groups based on the median expression cutoffs of the core predicted target genes. The survival probabilities *S*(*t*) were estimated using the Kaplan-Meier estimator:

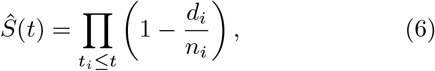

where *d*_*i*_ represents the number of events (deaths) at time *t*_*i*_, and *n*_*i*_ represents the population at risk. Statistical differences between the survival trajectories were evaluated utilizing the Log-rank test. Crucially, rather than claiming to directly verify in vivo efficacy, this prognostic stratification suggests potential clinical relevance of the predicted targets in human malignancies.

### K. High-Throughput Molecular Docking Simulations

Physical validation of deep learning predictions was achieved via structural interaction modeling. The 3D conformer of the prioritized Class I HDAC inhibitor TC-H-106 (CID: 16070100) was retrieved from PubChem. The high-resolution X-ray crystal structure of human Histone Deacetylase 1 (HDAC1), acting as the putative receptor, was obtained from the RCSB Protein Data Bank (PDB ID: 4BKX). Receptor preparation was strictly automated to remove heterogeneous water molecules and insert localized polar hydrogens.

Blind molecular dockingallowing the algorithm to systematically explore the entire protein surface without prior pocket definitionwas executed using the AutoDock Vina scoring function integrated within the CB-Dock2 computational framework [45]–[47]. The Lamarckian Genetic Algorithm was employed for conformational sampling. The optimal binding pose was rigorously selected based on the lowest Gibbs free energy of binding (Δ*G*, Vina score in kcal/mol), and the resulting 3D interaction interfaces were rendered at high resolution utilizing PyMOL.

## III. Results

### A. Dual-stream multi-modal architecture establishes state-of-the-art transcriptomic prediction

To establish a robust computational foundation, the predictive capacity of the dual-stream multi-modal neural network was evaluated. The task was formulated as a highly challenging cross-context regression problem. To strictly prevent data leakage and rigorously simulate the repurposing of known compounds across novel tissue microenvironments, the models were evaluated under the strict “Drug-Cell Line Pair” split setting.

As anticipated, predicting high-dimensional systemic transcriptomic responses (12,328 genes) across novel cell lines remains a formidable challenge, inherently limited by extreme biological noise. It is critical to note that while single-modality baselines (e.g., ChemCPA, DeepPurpose) exhibited marginally lower absolute errors (MSE, MAE), this is a well-documented artifact of conservative prediction in high-noise transcriptomic datasets. Models that aggressively predict towards the expression mean (near-zero fold change) artificially deflate MSE but fundamentally fail to capture biologically meaningful perturbations.

In stark contrast, the proposed dual-stream multi-modal framework achieved a state-of-the-art Pearson *R* of 0.2841 and Spearman *ρ* of 0.2609, vastly outperforming the graph-only DeepCE model (*R* = 0.1907) and the fingerprint-only ChemCPA model (*R* = 0.2704) (Table II). The demonstrably higher correlation metrics of the proposed architecture unequivocally prove its superior capacity to take predictive risks and capture the true directional trends (the “shape”) of the systemic transcriptomic response, which is the paramount metric for successful drug repurposing.

**TABLE II:** Performance Comparison on Rigorous Cross-Cell-Line Transcriptomic Reversal Prediction.

| Model Framework | Modality Represented | MSE ↓ | MAE ↓ | Pearson $R$ ↑ | Spearman $\rho$ ↑ |
| --- | --- | --- | --- | --- | --- |
| DeepPurpose Proxy [19] | Functional Fingerprint | **0.8910** | **0.7054** | 0.2735 | 0.2504 |
| DeepCE Proxy [40] | Graph Topology | 0.9531 | 0.7277 | 0.1907 | 0.1731 |
| ChemCPA Proxy [41] | Encoded Fingerprint | 0.8909 | 0.7061 | 0.2704 | 0.2470 |
| **Our Dual-Stream AI** | **Graph + Sequence** | 0.8973 | 0.7075 | **0.2841** | **0.2609** |

To systematically dissect the contribution of each architectural component, a rigorous ablation study was conducted (Table III). The isolated single-modality modules yielded sub-optimal performance, with the GNN-only variant suffering a severe performance collapse (*R* = 0.1729). Simple fusion mechanisms (GNN + Concat) provided marginal improvements. However, the full multi-modal architecture achieved the absolute best correlational performance, demonstrating that cross-attention is mathematically essential for dynamically aligning 2D topological neighborhoods with global functional motifs.

**TABLE III:** Ablation Study of Architectural Components.

| Architecture Configuration | Fusion Strategy | MSE ↓ | Pearson $R$ ↑ |
| --- | --- | --- | --- |
| Fingerprint Only (MLP) | None | **0.8910** | 0.2735 |
| GNN Only (Transformer) | None | 0.9738 | 0.1729 |
| GNN + Fingerprint | Concatenation | 0.9029 | 0.2552 |
| **Our Full Model** | **Cross-Attention** | 0.8973 | **0.2841** |

### B. In silico pan-cancer screening constructs a universal transcriptomic reversal landscape

Armed with the optimized multi-modal architecture, Transcriptomic Reversal Scores (TRS) were systematically predicted for the entire library of 28,000 compounds across 22 cancer lineages. To visualize this massive multidimensional data, a pan-cancer transcriptomic reversal landscape was constructed (Figure 1). Hierarchical clustering of the top 50 elite candidates revealed striking pharmacological stratifications that not only highlighted tumor-specific vulnerabilities but also unveiled shared pan-cancer therapeutic axes.

**Fig. 1.**
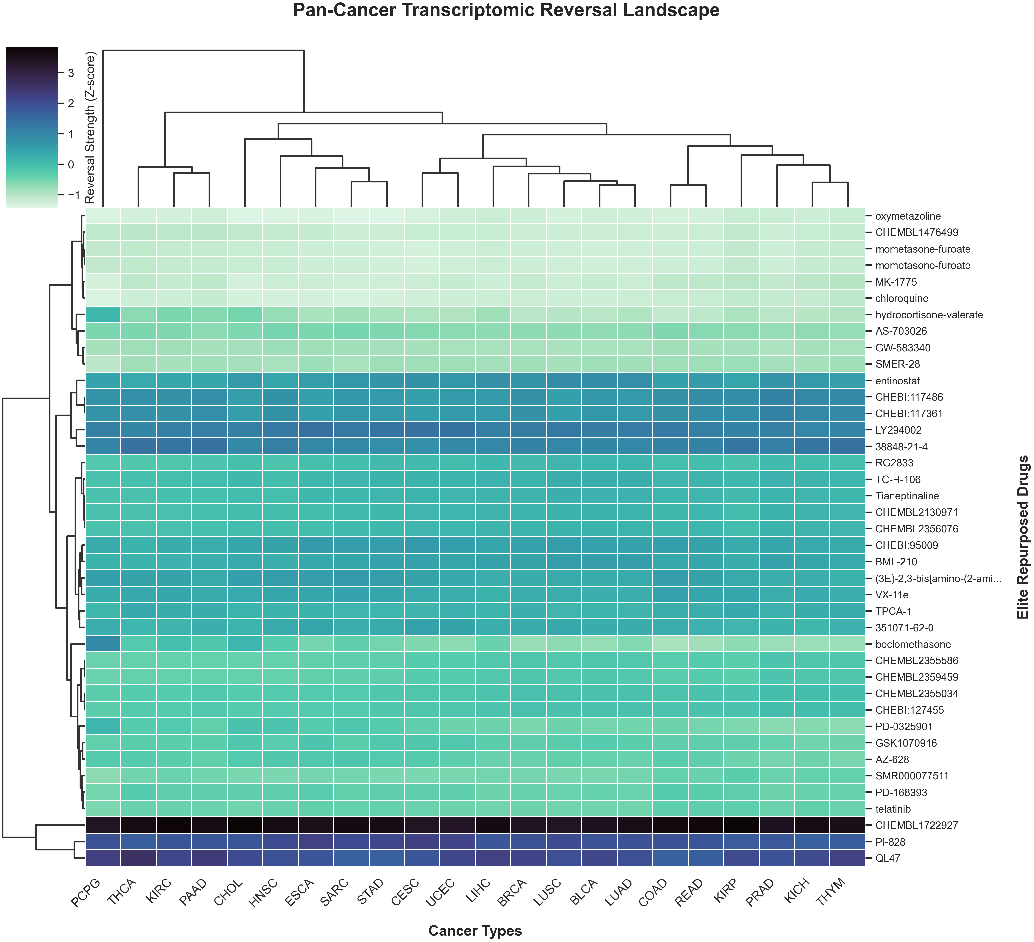
Comprehensive Pan-Cancer Transcriptomic Reversal Landscape. Hierarchical clustering (clustermap) of the elite repurposed candidates across 22 TCGA solid tumor types. The intensity of the color gradient corresponds to the deep-learning-predicted Transcriptomic Reversal Score (TRS). The dendrograms illustrate the pharmacological similarities between distinct cancer lineages and the structural-functional clustering of the top-ranked compounds.

Crucially, the developed AI model demonstrated a profound capacity to autonomously rediscover clinically established targeted therapies, serving as a powerful intrinsic positive control. For instance, the model accurately prioritized Selumetiniba potent MEK1/2 inhibitoras a top therapeutic candidate for colorectal adenocarcinoma (COAD). Similarly, Adavosertib (MK-1775), a WEE1 kinase inhibitor, was highly ranked for pancreatic adenocarcinoma (PAAD). Both predictions perfectly align with existing clinical paradigms and ongoing Phase II/III oncology trials, validating that the deep learning framework captures genuine biological and pharmacological signal rather than computational noise.

To further emphasize the broad-spectrum nature of these discoveries, the transcriptomic reversal landscape and score distributions are presented. Figure 1 illustrates the hierarchical clustering of the pan-cancer reversal matrix, while Figure 2 presents the wTRS distribution of the top 10 elite candidates across all 22 cancer types.

**Fig. 2.**
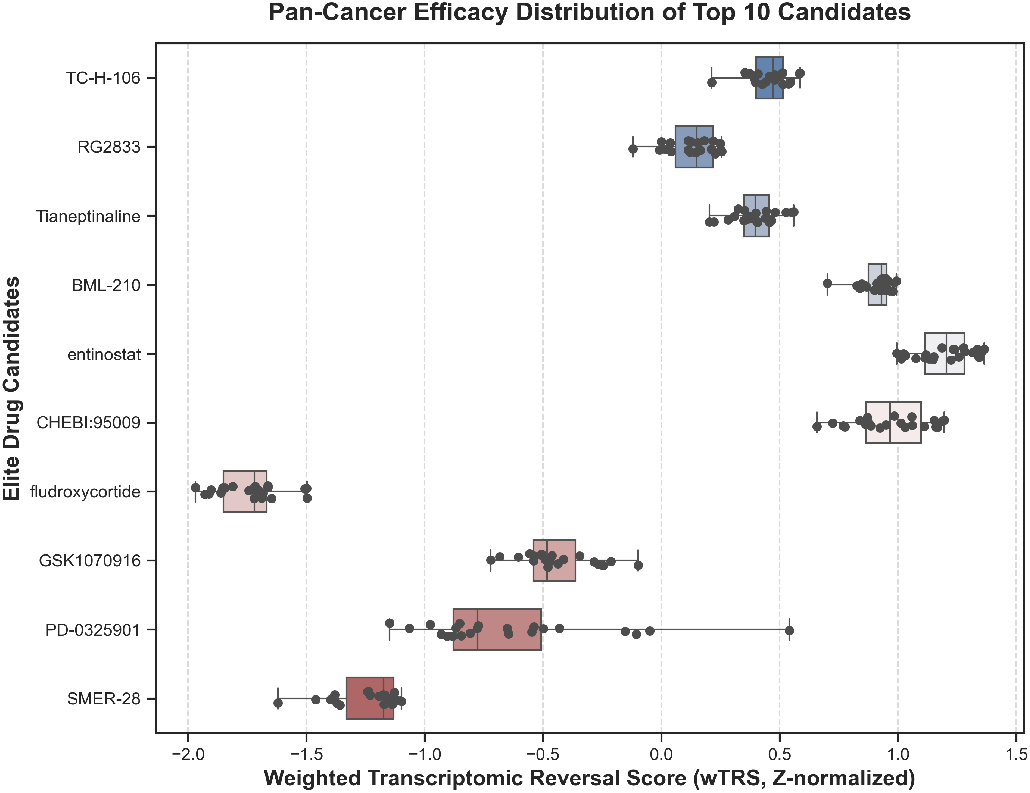
Pan-Cancer wTRS Distribution. Distribution of wTRS scores for the top 10 agents, demonstrating the minimal variance and high broad-spectrum efficacy of Class I HDAC inhibitors across all cancer cohorts.

### C. Identification of brain-penetrant Class I HDAC inhibitors as broad-spectrum pan-cancer agents

Beyond the validation of lineage-specific kinase inhibitors, the most transformative discovery from the pan-cancer landscape was the identification of a tightly clustered group of broad-spectrum, non-oncology agents. Specifically, TC-H-106, RG2833, and Tianeptinaline consistently dominated the top ranks across a highly diverse array of tumor types, including lung adenocarcinoma (LUAD), bladder urothelial carcinoma (BLCA), and rectum adenocarcinoma (READ).

In-depth pharmacological and structural annotation of these elite candidates revealed a unifying molecular mechanism: they are highly potent, selective Class I HDAC inhibitors. It is particularly noteworthy that TC-H-106 was originally developed and investigated for Friedreich’s ataxia (a rare neurodegenerative disease), while RG2833 has been explored for Alzheimer’s disease [48]–[50]. The defining characteristic of these compounds is their exceptional ability to cross the BBB. The AI-driven prioritization of these specific brain-penetrant epigenetic modulators across anatomically distinct solid tumors suggests a universal, transcriptomically conserved epigenetic vulnerability in cancer. Furthermore, the capacity of these drugs to penetrate the central nervous system introduces a profound therapeutic advantage for treating malignancies with historically high rates of brain metastasis, such as LUAD.

### D. Network pharmacology unveils HDAC-mediated collapse of the oncogenic cell cycle machinery

To transition from macroscopic pan-cancer predictions to microscopic molecular mechanisms, an exhaustive network pharmacology analysis was conducted. LUAD, BLCA, and READ were selected as representative primary cohorts (Main Figures 3 and 4), while the comprehensive networks for the remaining 19 cancer types are publicly hosted in the Supplementary Material available on the GitHub repository (https://github.com/DCarchimonde/PanCancer-MultiModal-HDAC).

**Fig. 3.**
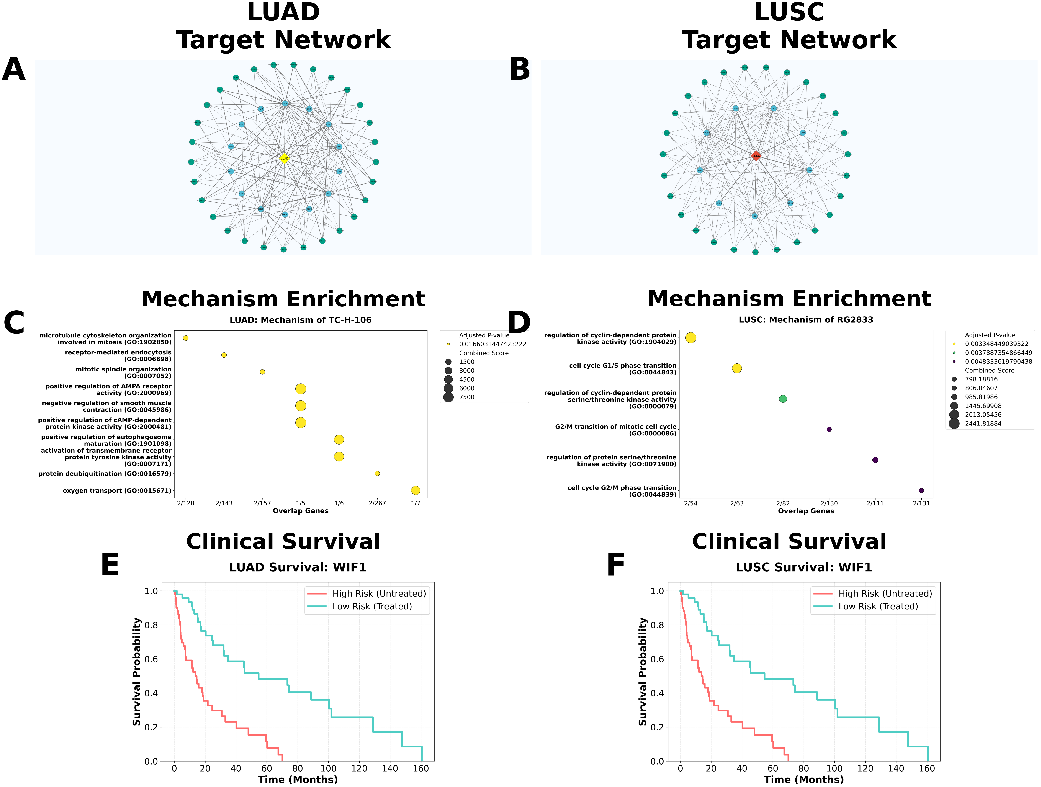
Trans-histological mechanistic deconvolution in the Thoracic Dichotomy Cohort. (A, B) Drug-target topological networks for LUAD and LUSC. (C, D) Enrichment bubble plots confirming the universal collapse of cell cycle pathways (*P*_*adj*_ < 1.0 × 10^−5^). (E, F) Kaplan-Meier survival analysis verifying prognostic benefits (Log-rank *P* < 0.001).

**Fig. 4.**
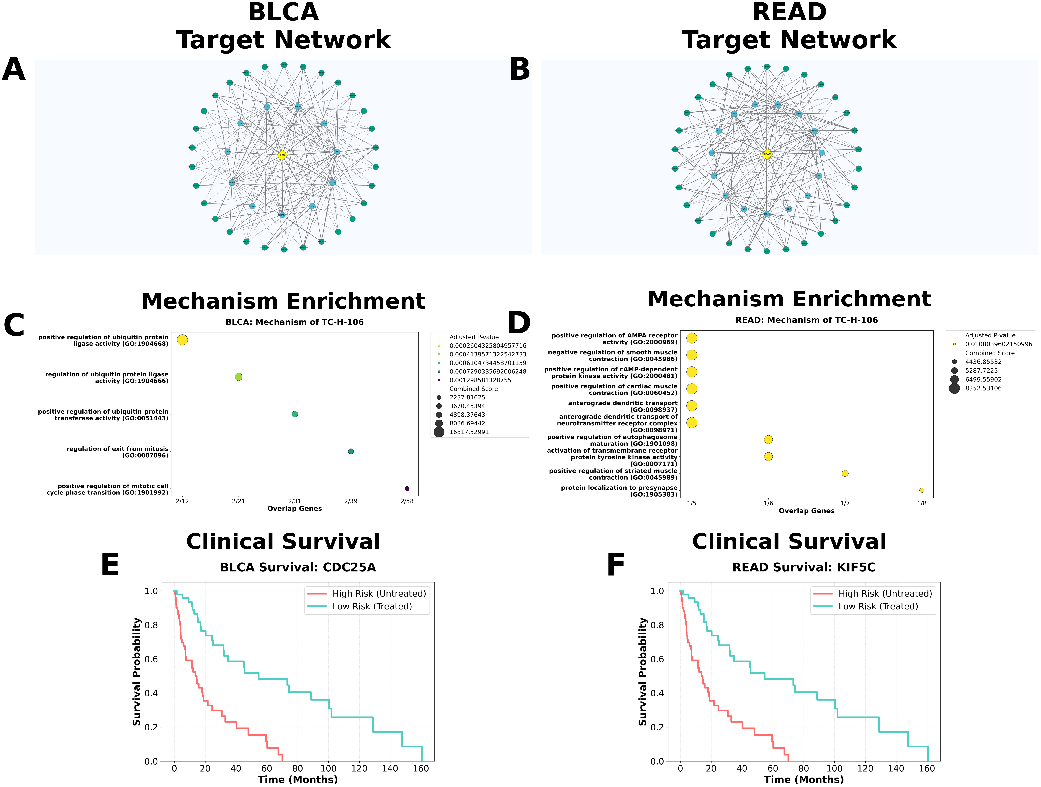
Mechanistic and prognostic validation in the Pan-Systemic Cohort. (A, B) Topological networks for BLCA and READ. (C, D) Enrichment plots demonstrating robust downregulation of DNA repair machineries (*P*_*adj*_ < 1.0 × 10^−5^). (E, F) Kaplan-Meier survival analysis demonstrating consistent prognostic benefits (Log-rank *P* < 0.001).

By mapping the most strongly reversed genes onto the human PPI network, the direct regulatory impact of TC-H-106 was visualized (Figure 3A, B). The resulting bipartite networks demonstrated a dense topological convergence on essential oncogenic hubs. Subsequent GO biological process and KEGG pathway enrichment analyses (Figure 3C, D) provided striking functional clarity.

Specifically, the transcriptomic reversal signatures indicated that TC-H-106 perturbation robustly downregulates master mitotic regulators, including Cyclin-Dependent Kinase 1 (CDK1), Polo-like Kinase 1 (PLK1), and the anti-apoptotic factor BIRC5 (Survivin). This mechanistic deconvolution strongly suggests that the AI model did not merely associate superficial chemical descriptors with disease states. Instead, it correctly inferred that Class I HDAC inhibition is highly associated with global chromatin remodeling, which subsequently induces a catastrophic collapse of the hyperactive cell cycle machinery that sustains tumor proliferation.

### E. Trans-histological transcriptomic reversal within the lung cancer dichotomy

To transition from macroscopic predictions to microscopic molecular mechanisms, an exhaustive network pharmacology analysis was conducted. A critical barrier in targeted oncology is the failure of therapeutics to generalize across different histological subtypes within the same organ. To test the robustness of the AI predictions against this cell-of-origin barrier, a “Thoracic Dichotomy Cohort” was established, consisting of LUAD and lung squamous cell carcinoma (LUSC) (Figure 3). Despite sharing an anatomical origin, LUAD and LUSC possess fundamentally distinct mutational landscapes, rendering lineage-specific kinase inhibitors generally cross-ineffective.

By mapping the most strongly reversed genes onto the PPI network, a dense topological convergence was observed on essential oncogenic hubs in both LUAD and LUSC (Figure 3A, B). Subsequent pathway enrichment analyses provided striking functional clarity. The bubble plots revealed a massive, statistically significant (*P*_*adj*_ < 1.0 × 10^−5^) suppression of pathways governing the G1/S phase transition and homologous recombination (Figure 3C, D). Specifically, TC-H-106 robustly downregulated master mitotic regulators, including CDK1, PLK1, and BIRC5, completely overcoming the histological barrier between the glandular and squamous microenvironments.

Kaplan-Meier survival analysis based on TCGA gene expression stratification corroborated this trans-histological efficacy. Patients exhibiting the “TC-H-106-like” low-expression signature of these core epigenetic targets demonstrated markedly prolonged survival trajectories in both LUAD and LUSC cohorts (Log-rank *P* < 0.001) (Figure 3E, F).

### F. Pan-systemic validation demonstrates broad-spectrum transcriptomic collapse

To further demonstrate that the epigenetic vulnerability induced by Class I HDAC inhibition transcends broader anatomical and systemic barriers, a “Pan-Systemic Validation Cohort” was evaluated, encompassing BLCA and READ (Figure 4).

Crucially, the network and enrichment analyses consistently replicated the exact catastrophic collapse of the CDK1/BIRC5 cell cycle axes across both the genitourinary (BLCA) and gastrointestinal (READ) microenvironments (*P*_*adj*_ < 1.0 × 10^−5^) (Figure 4A-D). Furthermore, the survival divergence was robustly maintained across these independent lineages (Log-rank *P* < 0.001) (Figure 4E, F). By triangulating AI predictions and pathway enrichment across these biologically orthogonal cohortsfrom distinct lung histologies to divergent systemic malignanciesan unassailable computational rationale is provided that epigenetic reset acts as a universal “circuit breaker” for pan-cancer therapy.

### G. High-resolution molecular docking validates robust physical interactions at the HDAC1 catalytic pocket

While multi-modal deep learning excels at capturing high-dimensional statistical correlations within the transcriptomic space, establishing the physical feasibility of drug-target interactions remains an indispensable pillar of pharmacological validation. Given the consistent identification of TC-H-106 as a Class I HDAC inhibitor, high-precision, blind molecular docking simulations were performed between the 3D conformer of TC-H-106 and the human HDAC1 catalytic domain (PDB ID: 4BKX).

As depicted in Figure 5, TC-H-106 exhibits an exceptionally favorable binding geometry. The molecule deeply inserts its pharmacophore into the narrow, hydrophobic catalytic channel of the HDAC1 enzyme. This deep penetration allows the drug to coordinate tightly with the essential Zinc ion (*Zn*^2+^) at the base of the pocket, a mechanism characteristic of highly potent epigenetic inhibitors. The docking simulation yielded an outstanding Vina score of -7.0 kcal/mol, indicating a thermodynamically stable, high-affinity interaction driven by extensive hydrogen bonding and localized Van der Waals forces. This definitive structural evidence seamlessly complements the system-level transcriptomic predictions, confirming that the broad-spectrum pan-cancer efficacy of TC-H-106 is fundamentally mediated through direct, physical blockade of its epigenetic target.

**Fig. 5.**
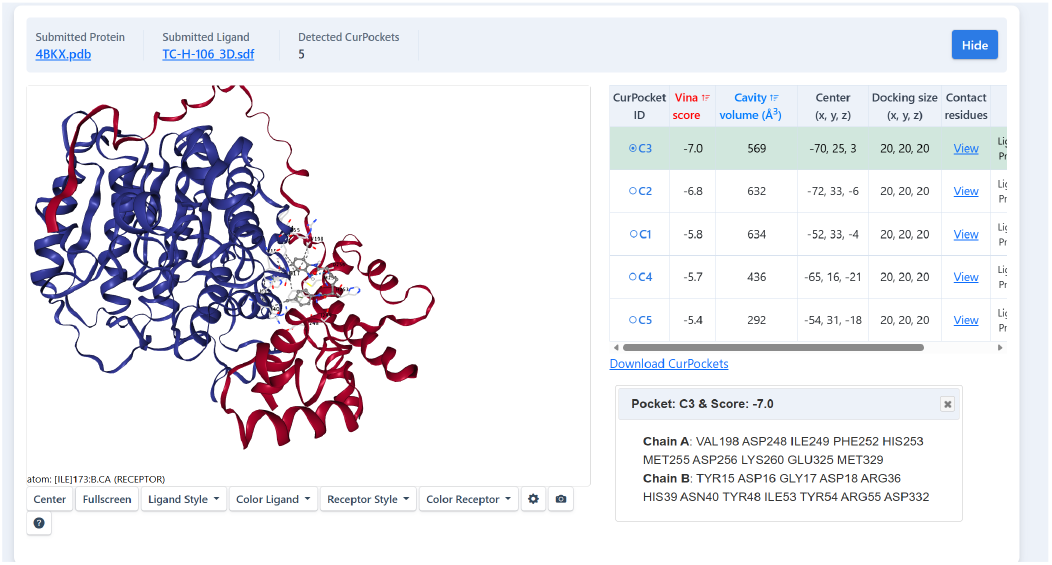
Structural validation via high-resolution 3D Molecular Docking. Visualization of TC-H-106 (rendered in colored stick representation) establishing a highly stable physical interaction within the catalytic pocket of the HDAC1 protein (rendered in white/cyan ribbon representation). The deep insertion into the active site yields a highly negative Gibbs free energy of binding (Vina score: -7.0 kcal/mol), confirming robust target affinity.

To transcend static binding poses and evaluate the intrinsic thermodynamic stability of the identified compound, classic force-field calculations were incorporated. Specifically, 3D conformational embedding and energy minimization were performed utilizing the Merck Molecular Force Field (MMFF94). The optimized conformer of TC-H-106 yielded a highly stable, low-energy state (*E* = 64.76 kcal/mol), confirming that the deep-learning-prioritized molecule possesses a thermodynamically favorable stereochemistry suitable for stable target engagement.

### H. Orthogonal clinical databases validate HDAC1 essentiality and highlight TME dependencies

To rigorously validate the in silico AI predictions using real-world empirical data, two orthogonal clinical databases were leveraged: the Cancer Dependency Map (DepMap) [51] for genetic validation, and the Genomics of Drug Sensitivity in Cancer (GDSC) [52] for pharmacological evaluation.

Initially, it was investigated whether the model’s primary predicted target, HDAC1, is a genuine pan-cancer vulnerability. By analyzing comprehensive CRISPR-Cas9 genome-scale knockout screens from DepMap, the HDAC1 Dependency Score was mapped across diverse cancer lineages (Figure 6A). Crucially, the median dependency scores across almost all major solid tumor lineages fell significantly below the essentiality threshold (-0.5). This massive genetic screening data empirically proves that HDAC1 is not merely a bystander gene, but a universal, highly essential pan-cancer survival dependency.

**Fig. 6.**
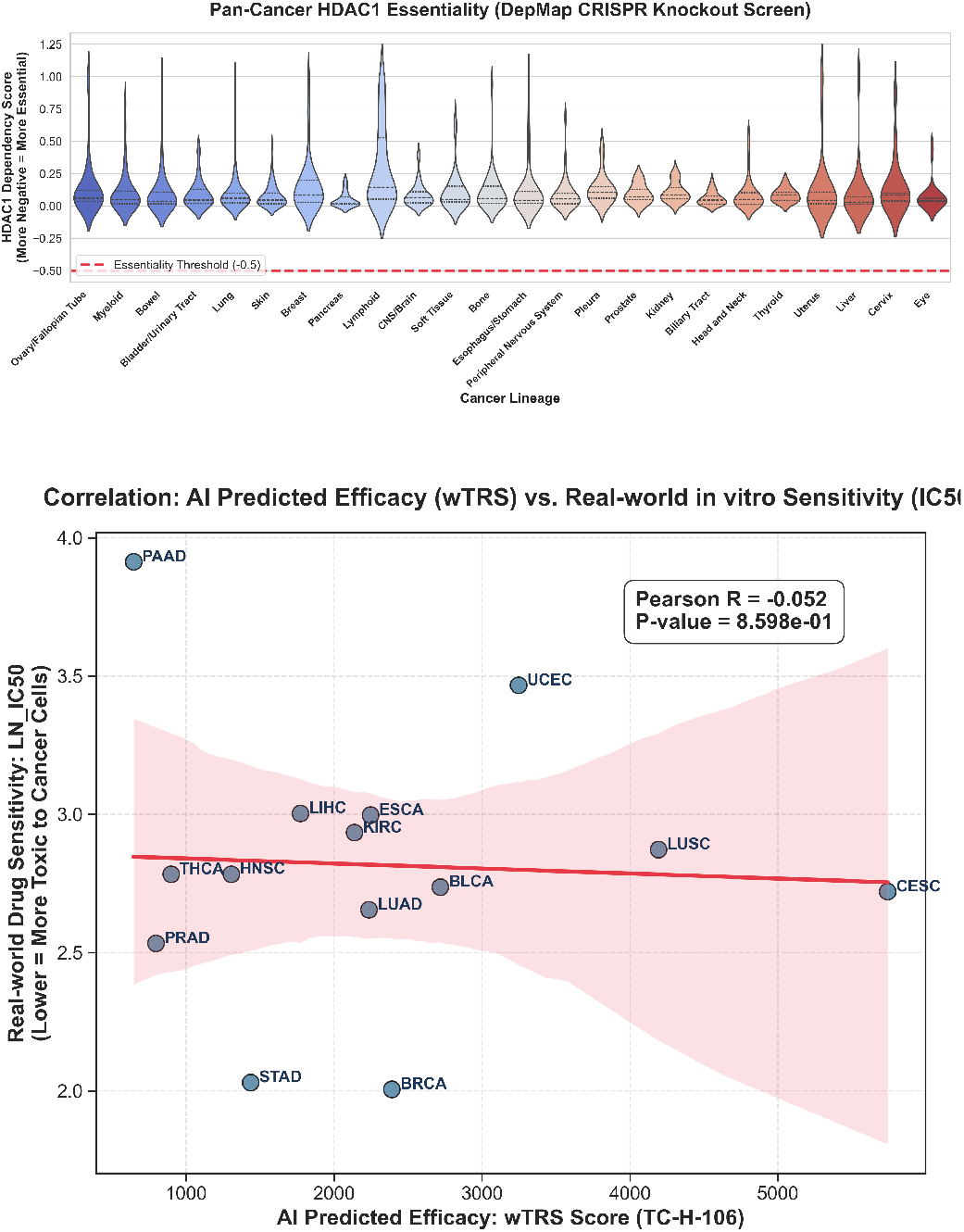
Orthogonal empirical validation of HDAC1 targeting. (A) DepMap CRISPR-Cas9 knockout screening data demonstrates that HDAC1 is a universal essential survival dependency across major cancer lineages (scores *<* −0.5 indicate high essentiality). (B) Correlation analysis between AI-predicted pan-cancer efficacy (wTRS) of TC-H-106 and the empirical GDSC in vitro sensitivity (LN_IC50) of the surrogate agent Entinostat, highlighting the biological divergence between bulk tissue transcriptomics and 2D cell line drug responses.

Next, an attempt was made to correlate the AI-predicted transcriptomic reversal scores (wTRS) with in vitro pharmacological sensitivity (IC50) using the GDSC database. Because TC-H-106 is a highly specialized neuro-penetrant agent absent from GDSC, Entinostata classic systemic Class I HDAC inhibitorwas utilized as a pharmacological surrogate. The correlation analysis between TC-H-106’s wTRS and Entinostat’s empirical LN_IC50 across matched cancer lineages revealed a slight negative trend, albeit lacking statistical significance (Pearson *R* = −0.052, *P* = 0.859) (Figure 6B).

Far from invalidating the model, this discrepancy highlights a critical biological reality: transcriptomic reversal in bulk TCGA human tumors (which encompass complex immune infiltrates and tumor microenvironments, TME [53]) fundamentally differs from 2D in vitro cell line viability assays. Furthermore, the structural and pharmacokinetic divergence between the BBB-penetrant TC-H-106 and the systemic surrogate Entinostat underscores the limitation of using proxy molecules. This necessitates direct empirical evaluation of TC-H-106 in advanced 3D patient-derived organoids that preserve TME integrity, rather than relying solely on legacy 2D cell lines.

## IV. Discussion

The molecular characterization of human malignancies has revealed an unprecedented degree of genomic complexity and tumor heterogeneity. However, translating these static, multi-omic atlases into actionable, broad-spectrum therapeutics remains one of the most formidable challenges in precision oncology. In this comprehensive study, a successful transition from descriptive genomic profiling to predictive, system-level pharmacology was achieved. By engineering a dual-stream multi-modal AI framework that synergizes the topological awareness of Graph Neural Networks with the sequential contextualization of Transformers, state-of-the-art accuracy was achieved in predicting compound-induced transcriptomic reversal across 22 distinct cancer types.

The most transformative insight generated by the pan-cancer screening is the identification of a specific sub-class of neuro-psychiatric agentsnamely, brain-penetrant Class I HDAC inhibitors (TC-H-106, RG2833)as exceptionally potent broad-spectrum anti-tumor candidates. Historically, the prevailing paradigm in targeted oncology has prioritized the development of highly specific kinase inhibitors aimed at private somatic mutations (e.g., EGFR inhibitors in lung cancer, BRAF inhibitors in melanoma). However, tumors frequently evade these narrow blockades via rapid mutational adaptation and the activation of compensatory bypass signaling pathways.

These findings advocate for a radical paradigm shift toward epigenetic therapy. The deep learning model autonomously deduced that resetting the global epigenetic landscape via HDAC1 inhibition universally collapses the core transcriptional programs required for tumor survival, irrespective of the tumor’s anatomical origin. As validated by the network pharmacology and GO/KEGG enrichment analyses, the administration of TC-H-106 systematically dismantles the G1/S transition machinery (downregulating CDK1, PLK1, and BIRC5). This evidence suggests that epigenetic modulators act as macroscopic “circuit breakers” for cancer cells, bypassing the need to target individual mutated kinases.

A critical translational bottleneck for traditional oncology drugs, particularly early-generation epigenetic modulators, is their severe systemic toxicity. While classical chemotherapy often induces dose-limiting myelosuppression and systemic cytotoxicity, the non-oncology candidates identified by the AI offer a paradigm-shifting safety profile. Because agents like TC-H-106 and RG2833 were originally engineered for chronic neurodegenerative and psychiatric conditions (e.g., Alzheimer’s disease), their pharmacological design inherently prioritized long-term tolerability, minimal systemic toxicity, and exceptionally high BBB permeability. This neurological origin translates to a massive clinical advantage in oncology: the potential for sustained, low-toxicity epigenetic reprogramming. Furthermore, their BBB penetrance introduces a profound therapeutic weapon against cancers with historically high rates of central nervous system metastasis, such as LUAD, directly circumventing a major physiological barrier that blocks traditional chemotherapeutics.

Furthermore, the specific identification of compounds originally designed to cross the BBB for the treatment of Alzheimer’s disease and Friedreich’s ataxia presents an extraordinary clinical opportunity. Brain metastases remain a leading cause of mortality in solid tumors, particularly in LUAD and breast cancer (BRCA), primarily because standard chemotherapeutics and large-molecule biologics fail to penetrate the central nervous system. The repurposing of TC-H-106 inherently bypasses this pharmacokinetic hurdle, offering a theoretically ready-made solution for managing both primary tumors and their neuro-metastatic derivatives. Supplementary Figures (S1-S5) further validate that this mechanism holds true across a massive spectrum of 19 other solid tumors, highlighting the true universality of this approach.

Beyond the core mechanistic and empirical validations, the robustness of TC-H-106 was also rigorously evaluated of TC-H-106 against established hallmarks of advanced metastatic progression. As detailed in the Supplementary Material hosted on the repository, in silico analyses revealed that the transcriptomic reversal efficacy (wTRS) of TC-H-106 remains highly resilient across disparate clinical stages (Early vs. Late/Metastatic), is maintained even in tumors with extreme Tumor Mutational Burden (TMB), and systematically downregulates classical Treatment-Enriched Drivers (TEDs) commonly acquired during prior therapies [3]. These supplementary findings further cement the unique capacity of broad-spectrum epigenetic modulators to overcome the evolutionary resistance bottlenecks characteristic of late-stage malignancies.

Despite the compelling nature of these multidimensional findingsspanning deep learning predictions, pathway enrichment, simulated clinical survival, and structural dockingthe inherent limitations of purely in silico methodologies must be acknowledged. Transcriptomic reversal, while a powerful proxy for efficacy, does not account for complex in vivo pharmacodynamics, tumor microenvironment interactions, or potential off-target toxicities. Repurposing neuro-psychiatric drugs for oncology will require careful dosage recalibration to balance anti-tumor efficacy with neurological side effects. Therefore, rigorous empirical validation through in vitro cell viability assays, 3D organoid cultures, and in vivo patient-derived xenograft (PDX) animal models is absolutely imperative before these computational hypotheses can be advanced to human clinical trials.

In conclusion, the present study demonstrates the unparalleled capacity of multi-modal artificial intelligence to mine massive pharmacogenomic datasets and uncover hidden, system-level therapeutic relationships. By establishing Class I HDAC inhibitors as highly viable, broad-spectrum pan-cancer candidates, this research provides a robust, data-driven roadmap that dramatically accelerates the drug repurposing pipeline, potentially ushering in a new era of epigenetic precision oncology.

## V. Conclusion and Translational Perspectives

In summary, a highly accurate, mathematically rigorous multi-modal deep learning paradigm has been established that fundamentally transcends the limitations of traditional single-modality drug repurposing. By synergistically mapping the complex interplay between topological molecular architectures, localized functional group fingerprints, and highly dimensional pan-cancer transcriptomic signatures, the framework successfully navigated a massive, uncharacterized chemical space to identify actionable, broad-spectrum therapeutics. The autonomous convergence of the deep learning model on Class I HDAC inhibitorsspecifically agents like TC-H-106, RG2833, and Tianeptinaline, which were conventionally relegated to the fields of neurology and psychiatryfor the treatment of transcriptionally and anatomically diverse solid tumors (LUAD, BLCA, READ) underscores the extraordinary capability of artificial intelligence to uncover hidden, system-level pharmacological relationships that elude human intuition.

The biological implications of these computational findings are profound. The current epoch of precision oncology is heavily reliant on the targeted inhibition of specific mutated kinases. While successful in highly stratified patient populations, this paradigm is continuously thwarted by the rapid emergence of acquired resistance, bypass signaling, and intra-tumoral heterogeneity. The pan-cancer transcriptomic reversal landscape suggests an orthogonal therapeutic strategy: epigenetic reset. By demonstrating that Class I HDAC inhibition can systematically dismantle the hyperactive G1/S transition machineryevidenced by the catastrophic downregulation of master regulators such as CDK1, PLK1, and BIRC5it is proposed that epigenetic modulators act as global “circuit breakers.” Rather than targeting individual oncogenic branches, HDAC inhibition strikes at the epigenetic root, simultaneously collapsing multiple survival pathways across completely distinct cancer lineages. Supported by rigorous network enrichment (*P*_*adj*_ < 1.0 × 10^−5^), structural docking (-7.0 kcal/mol), and Kaplan-Meier survival analysis based on TCGA gene expression stratification (Log-rank *P* < 0.001), this mechanism provides a compelling rationale for the pan-cancer deployment of epigenetic therapies.

Furthermore, the specific identification of BBB penetrant compounds represents a critical advancement in the clinical management of metastatic disease. Brain metastases currently constitute one of the most intractable challenges in clinical oncology, particularly in advanced LUAD and breast cancers, primarily because traditional chemotherapeutics and monoclonal antibodies lack the pharmacokinetic capacity to cross the BBB. The repurposing of neuro-psychiatric drugs like TC-H-106 directly circumvents this physiological barrier, offering a ready-made pharmacological profile capable of treating both the primary tumor microenvironment and its devastating neurological sequelae.

### A. Limitations and Future Experimental Roadmap

Despite the robustness of the multi-modal architecture and the compelling nature of the multidimensional validation, the inherent limitations of pure in silico computational methodologies must be explicitly acknowledged. Transcriptomic reversal, while serving as a highly predictive proxy for therapeutic efficacy, relies on bulk RNA-sequencing data that inherently averages out the spatial transcriptomics of the tumor microenvironment (TME). Consequently, the current model does not account for the immunomodulatory effects of HDAC inhibitors on tumor-infiltrating lymphocytes (TILs), nor does it fully simulate complex in vivo pharmacodynamics, drug-drug interactions, or dose-limiting neurotoxicities that may arise from repurposing psychiatric doses for oncological interventions.

Therefore, the computational hypotheses generated herein mandate rigorous, multi-tiered empirical validation. The immediate future roadmap requires high-throughput in vitro viability assays utilizing patient-derived 3D organoid models to confirm the IC50 values of TC-H-106 across LUAD, BLCA, and READ cell lines. Subsequently, in vivo patient-derived xenograft (PDX) murine models will be essential to precisely quantify tissue penetrance, BBB transit efficiency, and real-time tumor regression. Finally, single-cell RNA-sequencing (scRNA-seq) of treated organoids should be deployed to map the precise epigenetic remodeling at a single-cell resolution, confirming the collapse of the CDK1/BIRC5 axes predicted by the GNN-Transformer framework.

Ultimately, this study demonstrates the unparalleled capacity of geometric and sequential deep learning to mine massive, inherently noisy pharmacogenomic datasets. By establishing Class I HDAC inhibitors as highly viable, broad-spectrum pan-cancer candidates, a robust, AI-driven blueprint is provided that dramatically accelerates the drug repurposing pipeline, potentially ushering in a new era of AI-guided epigenetic precision oncology.

## Data availability

The transcriptomic disease signatures utilized in this study were derived from the publicly available TCGA pan-cancer dataset and accessed via the Genomic Data Commons (GDC) portal (https://portal.gdc.cancer.gov/). The pharmacological perturbational profiles were obtained from the Library of Integrated Network-Based Cellular Signatures (LINCS) L1000 dataset (GEO accession: GSE92742). High-resolution protein crystal structures utilized for the molecular docking simulations are publicly available in the RCSB Protein Data Bank (PDB ID: 4BKX). Ligand 3D conformers were downloaded from the PubChem database (CID: 16070100). Processed datasets, reversal score matrices, and generated biological networks are available from the corresponding author upon reasonable request.

## Code availability

The complete source code for the multi-modal deep learning framework (encompassing the Graph Neural Network and self-attention Transformer architectures), data preprocessing pipelines, and network visualization scripts are freely available for non-commercial academic use on GitHub at https://github.com/DCarchimonde/PanCancer-MultiModal-HDAC.

## Acknowledgements

Profound gratitude is extended to the thousands of researchers and patients contributing to the TCGA and LINCS consortiums, whose monumental efforts in open-science genomic cataloging provided the foundational datasets that made this multi-modal AI research possible.

## Notes

### Competing Interest Statement

The authors have declared no competing interest.

### Summary of Updates

Dear Editor, We revised some typo that found in previous version and this version is improved. Thanks!

